# A deep genetic structure phylogenomically frames the closest algal relatives of land plants

**DOI:** 10.64898/2026.04.20.719584

**Authors:** Maaike J. Bierenbroodspot, Cäcilia F. Kunz, Elisa S. Goldbecker, Maike Lorenz, Iker Irisarri, Thomas Pröschold, Tatyana Darienko, Jan de Vries

**Affiliations:** University of Göttingen, Institute for Microbiology and Genetics, Department of Applied Bioinformatics, Goldschmidtstr. 1, 37077 Göttingen, Germany; University of Göttingen, Albrecht-von-Haller-Institute for Plant Sciences, Department of Experimental Phycology and SAG Culture Collection of Algae, Nikolausberger Weg 18, 37073 Göttingen, Germany; Museo Nacional de Ciencias Naturales (MNCN-CSIC), Department of Biodiversity and Evolutionary Biology, José Gutiérrez Abascal 2, 28006 Madrid, Spain; University of Innsbruck, Research Department for Limnology, Mondseestr. 9, 5310 Mondsee, Austria; University of Göttingen, Campus Institute Data Science (CIDAS), Goldschmidtstr. 1, 37077 Göttingen, Germany; University of Göttingen, Göttingen Center for Molecular Biosciences (GZMB), Department of Applied Bioinformatics, Goldschmidtstr. 1, 37077 Göttingen, Germany; University of Maastricht, Maastricht Centre for Systems Biology and Bioinformatics (MaCSBio), Paul-Henri Spaaklaan 1, 6229 EN Maastricht, The Netherlands

**Author notes:** contributed equally (Tatyana Darienko, Jan de Vries). Correspondence (J.d.V.).

## Abstract

To discern the nature of the closest streptophyte algal relatives of land plants (embryophytes) is a major question in the field of plant evolutionary biology; discerning that nature is essential for our ability to infer the last common ancestor of embryophytes and algae, allowing to retrace the adaptations that ushered in the conquest of land by plants. Albeit initially coming as a surprise, all major phylogenomic efforts have concluded that the Zygnematophyceae are the algal sisters to land plants^1-6^. The Zygnematophyceae are the streptophyte algal class with the greatest species richness^7^ and while we now have ample genome information on a few select members of zygnematophytes^8-13^, our understanding of the genetic and genomic divergence as well as potential is limited by a lack of phylodiverse data that accounts for this diversity and integrates it into a phylogenomic framework. We here sequenced 43 new transcriptome datasets for the Zygnematophyceae and built a phylogenomic tree based on a total of 104 zygnematophyceaen transcriptomes and 2243 loci. We recover a deep genetic structure for the Zygnematophyceae, revealing that this algal class is ancient. Despite the deep split between Spirogyrales and their unicellular sister group Desmidiales, most Spirogyrales emerged after pronounced genetic divergence, accommodating the attainment of multicellularity and divergent traits such as unique cell and plastid division^13^. Overall, our data capture signatures of massive ancient radiations. Zygnematophyceae are characterized by deep genetic divergences that necessitated a phylodiverse sampling to be revealed, together with their vast evolutionary history, and to illuminate the nature of the algal progenitors of land plants.

## Main

As the closest algal relatives of land plants, Zygnematophyceae have attracted considerable attention from the plant evo-devo community in recent years^14-16^. The class Zygnematophyceae was established by Round^17^ and validated^18^. Traditionally, the class was divided into two orders, the filamenteous Zygnematales and the unicellular Desmidiales, based on morphology of vegetative cells^18,19^ Recently, based on a phylogenomic tree of 326 loci, Zygnematophyceae were divided into a five-order system^7^. While each order was well-supported, the phylogenetic relationship among them remained open and orders were in some cases represented by low genetic diversity. For this, we used a systematic sampling approach, aiming for covering maximized phylodiversity in Zygnematophyceae across taxonomic groups, growth types, habitats, and culture collections.

We obtained 43 strains from which we generated *de novo* transcriptomes. Combined with publicly available data, we assembled a dataset of 104 zygnematophyceaen algae, nine other streptophyte algae, eight land plants, and four chlorophyte algae as outgroup. Building on our previously established methodological approaches^20,21^ and IQ-Tree v3 we computed a phylogenomic tree based on 2243 loci and the site-heterogeneous best-fit model^22^ with Posterior Mean Site Frequency (PMSF)^23^ profiling (LG+C60+F+G-PMSF) (Figure 1).

**Figure 1:**
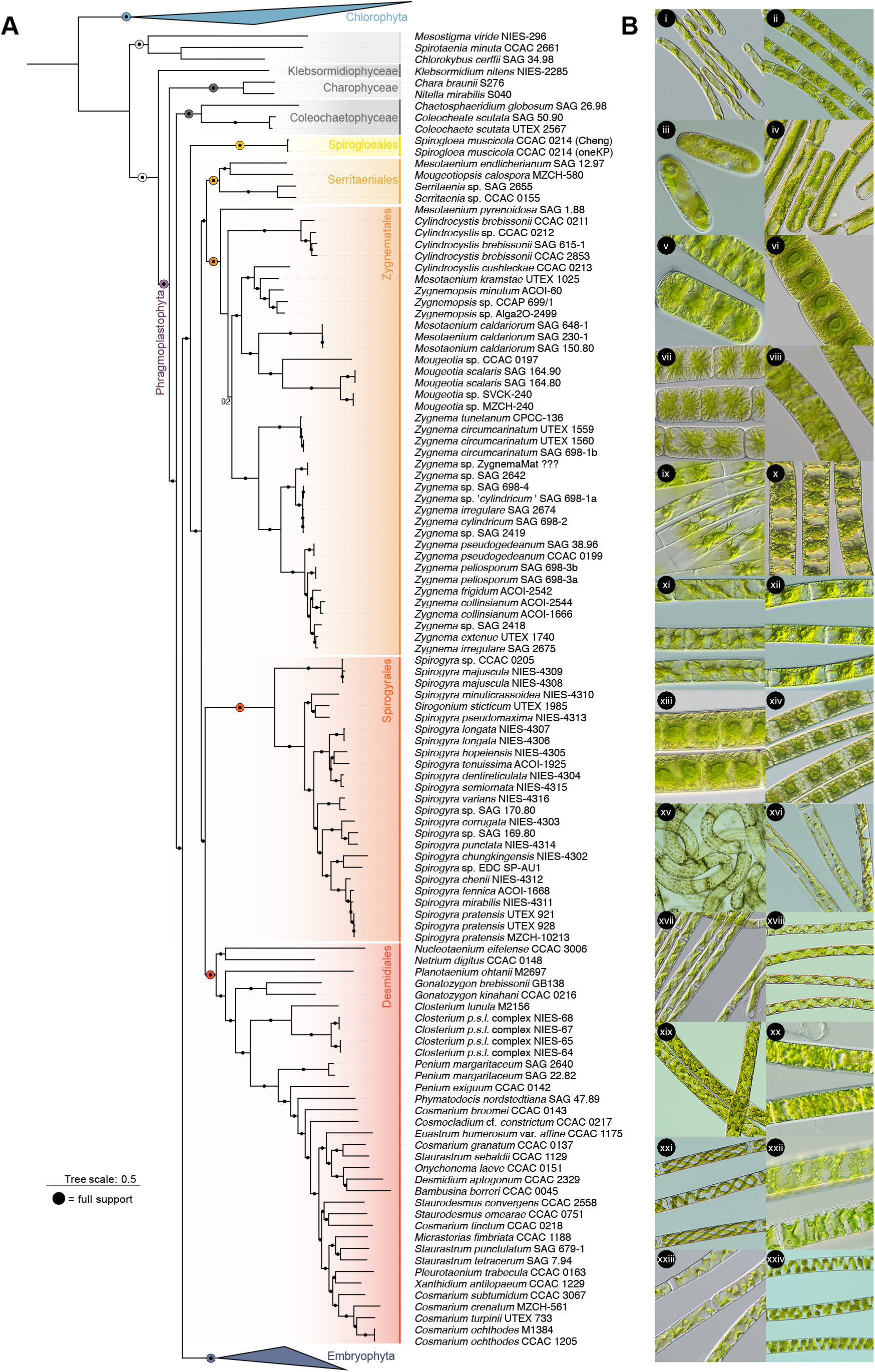
A robust phylogenomic framework and deep genetic structure within Zygnematophyceae. **(A)** A maximum likelihood phylogenomic tree based on 2243 loci. The Zygnematophyceae are divided into five orders: (i) Spirogloeales, (ii) Serritaeniales, (iii) Zygnematales, (iv) Spirogyrales, and (v) Desmidiales. Filled dots mark full support and the colors correspond to the labels of major monophyla. **(B)** Light micrographs showcasing the here newly sequenced diversity: i. *Mesotaenium kramstae* NIES-657; ii. *Zygnemopsis* sp. ACOI-60; iii. *Mesotaenium caldariorum* SAG 230-1; iv. *Mougeotia scalaris* SAG 164.80; v. *Zygnema circumcarinatum* 698-1b; vi. *Zygnema* sp. SAG 698-4; vii. *Zygnema cf. cylindricum* SAG 698-1a; viii. *Zygnema cylindricum* SAG 698-2; ix. *Zygnema pseudogedeanum* SAG 38.96; x. *Zygnema peleosporum* SAG 698-3a; xi. *Zygnema frigidum* ACOI-2542; xii. *Zygnema colinsianum* ACOI-1666; xiii. *Zygnema* sp. SAG 2418; xiv. *Zygnema* sp. SAG 2675; xv. *Spirogyra majuscula* NIES-4308; xvi. *Sirogonium sticticum* UTEX 1985; xvii. *Spirogyra tenuissima* ACOI-1925; xviii. *Spirogyra dentireticulata* NIES-4304; xix. *Spirogyra corrugata* NIES-4303; xx. *Spirogyra* sp. SAG 170.80; xxi. *Spirogyra chunkingensis* NIES-4302; xxii. *Spirogyra* sp. SAG 169.80; xxiii. *Spirogyra fennica* ACOI-1668; xxiv. *Spirogyra pratensis* MZCH-10213.

### The phylogenomic frame of Zygnematophyceae is robust

Our tree recovered the well-accepted gross topology of the Chloroplastida, with deeply split monophyletic chlorophytes and streptophytes, six monophyletic streptophyte algal classes, and the zygnematophytes as sister to land plants^1-5^ (Figure 1). Our sampling effort focussed on increasing the representation especially for the orders Spirogyrales and Zygnematales to improve the phylogenomic resolution of our analyses.

The traditional order Zygnematales was split into three orders as in ref.^7^: Spirogyrales, Serritaeniales, and Zygnematales (in revised form), which included the transfer of unicellular taxa (*Mesotaenium* Naegeli, *Cylindrocystis* Meneghini ex De Bary) to these new orders. Most of the other unicellular taxa remained in the Desmidiales. Interestingly, the fifth order Spirogloeales, remains represented by a single species, *Spirogloea muscicola* (De Bary) Melkonian, which was originally assigned as *Spirotaenia bryophila* (Brébisson) Lütkemüller ^8^. Overall, our phylogenomic study confirmed the five-order system^7^, but Zygnematales and Serritaeniales were recovered as sister groups in our analyses based on increased sampling of taxa and genes. Additional sampling of taxa such as members of the genera *Ancylonema* Berggren, *Zygogonium* Kützing, and *Mesotaenium-*like organisms, which are closely related based on preliminary *rbc*L phylogeny and genomic signatures^24-26^, could further change the topology of the orders Zygnematales and Serritaeniales and/or modify the orders as currently defined.

Building on the phylogenomic tree, we performed ancestral character state reconstruction (ACSR) to reconstruct the evolution of growth types across Zygnematophyceae (Figure 2). Based on the current sampling, of the seven transitions two filamentous growth, only two are ancient: in the last common ancestor (LCA) of Spirogyrales and the genus *Zygnema*—to both of which we turn in the following.

**Figure 2:**
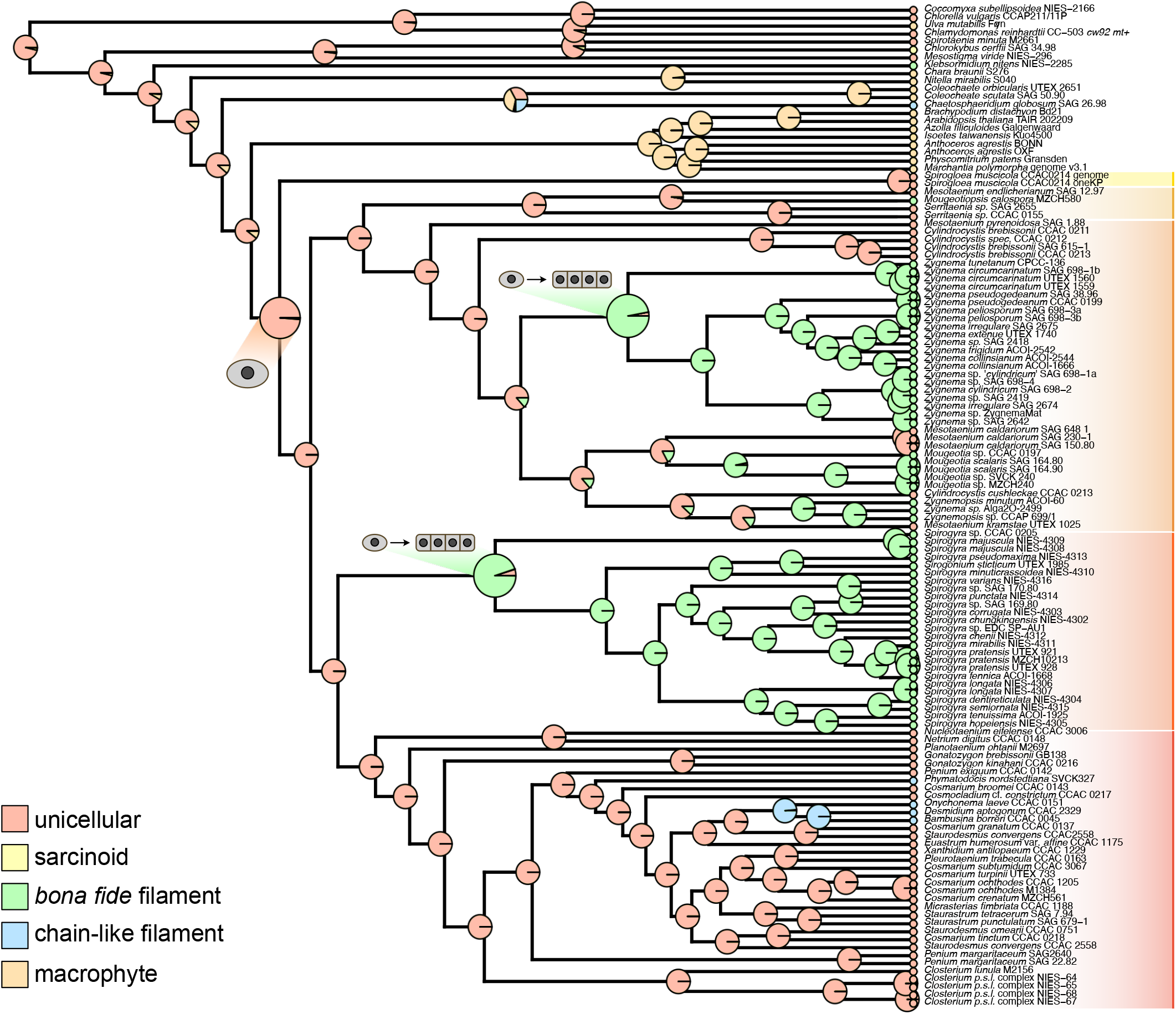
Ancestral character state reconstruction of body plan evolution across Zygnematophyceae. Growth types were coded as five-character state distributions to trace the evolution of different homologies of growth types: salmon = unicellular, yellow = sarcinoid, green = bona fide filamentous growth, blue = chain-like filaments (desmids), and orange = complex multicellular macrophytes. The inferred ancestral unicellular state for the LCA of all Zygnematophyceae is highlighted as well as the two transitions to filamentous growth types lead to the LCAs of the genus *Zygnema* and the order Spirogyrales.

### Pronounced ancient divergence before the emergence of extant Spirogyrales

The order Spirogyrales currently consists of the single genus *Spirogyra* Link. Its most iconic feature is its spiral chloroplast(s), undulating through the entire length of the cell. From the almost 400 species described, the genome of one species (*Spirogyra pratensis* Transeau) has been sequenced^13^. This single genome^13^—bolstered by the confirmation of patterns from previous draft sequencing data^27^—uncovered conspicuous deviations from other zygnematophytes, evident by genomic streamlining and pronounced genetic divergence; the most palpable example being the absence of the plastid division machinery otherwise conserved across Archaeplastida^13^. We thus used our phylogenomic framework to contextualize this divergence.

The only reliably way of distinguishing the different *Spirogyra* species based on morphology is by observing the formation of zygotes (zygospore) and their characteristics^28^. Unfortunately, the trigger for the induction of the sexual reproduction is unknown. Therefore, most of the available *Spirogyra* strains are unidentified at the species level. Takano et al.^29^ identified 13 species based on experiments inducing sexual reproduction. They sequenced two plastid-coding markers (*rbc*L and *atp*B) and discovered seven clades (I-VII). Our phylogenomic tree recovered seven highly supported clades of *Spirogyra* spp., which correspond to those established by Takano et al. (2019) with one exception: clade VI is missing because no strain of this clade is available in public culture collections. A subdivision into eight clades was found using SSU rDNA sequences (clades A-H; Chen et al.^30^). Unfortunately, no strain of this study is available and almost none was assigned to species because of the lack of zygospores in culture. Both of these studies confirmed that the genus *Sirogonium* Kützing 1843 belonged to *Spirogyra* (clade II in Takano et al., 2019 or clade C in Chen et al., 2012). This genus differed from *Spirogyra* by 2-10 ribbon-like chloroplasts, a feature which derives from spiral chloroplast(s) based on phylogenetic and phylogenomic studies. In addition, one new clade could be discovered in our phylogenomic study containing three strains (NIES-4303 *Sp. corrugata* Transeau, NIES-4314 *Sp. punctata* Cleve and SAG 169.80 *Spirogyra* sp.). Both NIES strains were not assigned to any clade in Takano et al. (2019).

Overall, our phylogenomic tree included a phylodiversity of 25 different *Spirogyra* species. They form a monophylum, the order Spirogyrales, that branches off from the sister order Desmidiales. The branch leading to extant Spirogyrales is remarkably long (6.2 longer than the branch leading to Desmidiales), which could be indicative of massive extinction or unsampled Spirogyrales diversity (currently undescribed). This would mean that emergence of signature traits such as the spiral chloroplasts, genomic streamlining, and unique division machineries co-changing during evolution, coming to bear before the LCA of extant Spirogyrales. This aligns with the ACSR, recovering ancient filamentous growth forms for Spirogyrales (Figure 2).

### Deep genetic structure and recent bursts of radiations in the genus *Zygnema*

The genus *Zygnema* C. Agardh is characterized by two asteroid chloroplasts, each with a single central pyrenoid; we recover them as ancestrally filamentous (Figure 2). 137 species are described based on features of zygospores^28^. Phylogenetic analyses of *rbc*L sequences revealed that all investigated specimens were distributed into three clades^31,32^. Our phylogenomic analyses confirmed this subdivision. Three groups could be discovered: *Z. circumcarinatum*-, *Z. cylindricum*- and *Z. peliosporum*-groups. The first clade contains only two species *Z. circumcarinatum* Czurda (SAG 698-1b) and *Z. tunetanum* (Gauthier-Lièvre) Stancheva, J.D.Hall, McCourt & Sheath (CPCC 136) the latter originally assigned to the genus *Zygogonium*. The second clade consists of several species, some not assigned at the species level. Interestingly, the strain SAG 698-1a, originally assigned as other mating type of *Z. circumcarinatum* (now *Z*. sp. ‘*cylindricum*’) belongs to this clade, which indicated a past mistransfer in algal collections^10^. Most of the investigated strains are members of the third clade.

The sister group of Zygnematales in our phylogenomic study is the order Serritaeniales^7^. This order includes (i) several taxa assigned to the genus *Serritaenia* A.Busch & S.Hess, (ii) the type species of the genus *Mesotaenium, M. endlicherianum* Naegeli, and (iii) *Mougeotiopsis calospora* Palla. Interestingly, *Mougeotiopsis* represents the only filamentous taxon within this order so far. The other specimens of this order have unicellular *Mesotaenium*-like morphologies. The genus *Mesotaenium* is polyphyletic as currently described. For example, *M. pyrenoidosum* (P.A.Broady) Petlovany and *M. kramstae* Lemmermann belong to different lineages of the Zygnematales. The diversity of *Mesotaenium*-like organisms is much higher than expected and await taxonomic revisions^24^.

Our phylogenomic data recovers the genus *Zygnema* as monophyletic. In previous *rbc*L phylogenies^24^, *Zygnema* was placed in an unsupported larger group consisting of other species of *Cylindrocystis*, different lineages of *Mesotaenium* (Meso-9 and Meso-10), and *Zygnemopsis* (Skuja) Transeau. Our phylogenomic study now robustly resolves their relationships; three additional, well-supported lineages beside the *Zygnema* subclade are recovered: (i) *Zygnemopsis* together with *Cylindrocystis cushleckae* A.J.Brook and *Mesotaenium kramstae*, (ii) *Mougeotia* C.Agardh and *Mesotaenium caldariorum* (Lagerheim) Hansgirg (Meso-9), (iii) *Cylindrocystis brebissonii*, (iv) and *Mesotaenium pyrenoidosum* (Meso-4) sister to all other Zygnematales. Both genera *Zygnema* and *Zygnemopsis* cannot be told apart by looking at the vegetative filaments, but by the presence of mucilage surrounding the gametangia in *Zygnemopsis*. Compared to *Zygnema* and *Spirogyra, Zygnemopsis* is a small genus consisting of 42 species^28^. In our analyses, the investigated strains are separated from *Zygnema* confirming that this morphological feature seems apt for differentiation of both genera if sexual stages are present. Unfortunately, no sexual reproduction was observed for the isolates included in the tree and therefore, no species assignment was possible.

Based on the topology of our phylogenomic tree we made two observations for the extant diversity represented by the genus *Zygnema*. First, the sub-clades within the genus are deeply divergent from one another—aligning with the molecular clock analysis of Feng et al.^10^ that dated the divergence between *Zygnema circumcarinatum* and *Zygnema cylindricum* Transeau to have occurred 236±5 million years ago. Second, each sub-clade experienced pronounced radiation, giving rise to the diversity captured by our dataset.

### Order-specific occurrence of signature genes shared by land plants and zygnematophytes

Traditionally, other streptophyte algal lineages than Zygnematophyceae were considered as closest algal relatives of land plants. This was owed to single gene phylogenies and the complexity of body plans found in genera like *Chara* L. and *Coleochaete* Brébisson; yet, both in the case of Charophyceae and the discoidal *Coloechaete* species the complexity in body plans are the result of secondary gains^21,33^. The genomic revolution has not only phylogenomically established the sister group relationship of Zygnematophyceae and embryophytes^1-3,34^ but comparative genomics pinpointed to several genetic synapomorphies in the ancestor of Zygnematophyceae and embryophytes^35^.

The genes homologous to those coding for the important canonical abscisic acid receptor^36-42^ PYRABACTIN RESISTANCE1/PYR1-LIKE/REGULATORY COMPONENTS OF ABA RECEPTOR (PYR/PYL/RCAR) are found in the genomes of *Mesotaenium*^8^ and *Zygnema*^10,35^ but not in that of the desmids *Penium* Brébisson ex Ralfs ^9^ and *Closterium* Nitzsch ex Ralfs ^11^. In light of this patchy recovery of signature genes shared by land plants and zygnematophytes, genome space of the Zygnematophyceae has been insufficiently covered only by one or two genomes per order. We here made use of our largely extended sampling to explore the occurrence of *PYR/PYL/RCAR* homologs in zygnematophytes. Our maximum likelihood phylogeny recovers that the well-known *PYL* homologs of *Mesotaenium endlicherianum* SAG 12.97 and *Zygnema circumcarinatum* SAG 698-1b fall into monophyla of *PYL* homologs of Serritaeniales and Zygnematales, respectively (Figure 3A)—including homologs in, e.g., *Zygnemopsis*, which is within the order Zygnematales and distantly related to the model strains of *Zygnema*. Yet, despite our vastly extended dataset, we did not detect homologs in any other order, suggesting likely order-specific losses of *PYL* (one in Spirogloeales and one in the common ancestor of Desmidiales and Spirogyrales) or the need for extended sampling.

**Figure 3:**
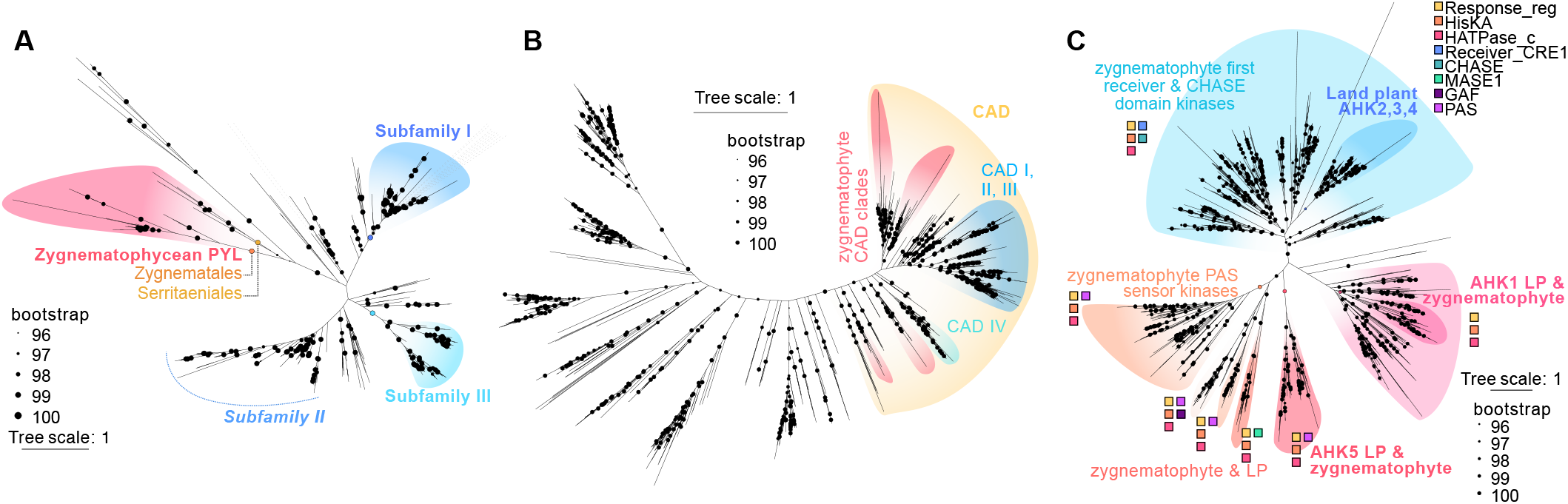
Order-specific radiations in land plant heritage genes. Maximum likelihood phylogenies of protein-coding genes homologous to signature regulators of the molecular biology of land plants. (A) Phylogeny of *PYRABACTIN RESISTANCE1/PYR1-LIKE/REGULATORY COMPONENTS OF ABA RECEPTOR* (*PYR/PYL/RCAR*) homologs, radiating independently of the land plants subfamilies (cf. Park et al.^40^ and Zimran et al.^83^) in Zygnematophyceae and limited to the orders Serritaeniales and Zygnematales. (B) Phylogeny of *CINNAMYL ALCOHOL DEHYDROGENASE* (*CAD*), showing several order-specific clades and radiations; homologs were recovered across orders. (C) Phylogeny of histidine kinase homologs and their domain structure. PFAM domains were annotated using interproscan and consolidated per clade. PF00072: Response regulator receiver domain (Response_reg), PF00512: His Kinase A (phosphor-acceptor) domain (HisKA), PF02518: Histidine kinase-, DNA gyrase B-, and HSP90-like ATPase (HATPase_c), PF03924: CHASE domain, PF24896: AHK4/CRE1/WOL first receiver domain (Receiver_CRE1), PF05231: MASE1, PF13185: GAF domain, PF00989: PAS fold (PAS), PF08447: PAS fold (PAS_3), PF08448: PAS fold (PAS_4), PF13188: PAS domain (PAS_8), PF13426: PAS domain (PAS_9). For the fully labelled trees, see Supplemental Figure S1.

Another set of land plant heritage genes concerns the chassis of the phenylpropanoid pathway. The phenylpropanoid pathway is key to a myriad of aspects of embryophyte biology, leading to the production of compounds crucial for biotic and abiotic stress response^43,44^ as well as structural compounds such as lignin^45,46^. Tracing its evolutionary origin to a chassis that emerged in algae is thus a long-standing question in the field^47-52^. Streptophyte algae not only have genes homologous to those coding for a series of enzymes that have assembled in land plants into the phenylpropanoid pathway, but in some cases these homologs have even a radiated repertoire^53^. *CINNAMYL ALCOHOL DEHYDROGENASE* (*CAD*) is a point in case^53^. CAD is the last step in monolignol biosynthesis, reducing diverse cinnamyl aldehydes into their corresponding cinnamyl alcohols^54,55^.

Homologs of this NADP(H)-dependent enzyme have been classified into three to five different groups or classes. Generally, *bona fide*^56^ or class I CADs show a clear response to stress^57-60^ with the highest catalytic activity^57,61,62^ and with the full catalytic motif set^63,64^. The other groups or classes describe CAD-like isoforms with generally a lower (Class II)^57,61,62,65^ to an even lower or no catalytic activity (Class III)^61,65^ or other functions. We here use the classification from Saballos et al.^66^, where group IV was described as monocot-specific.

None of the *CAD* homologs of Zygnematophyceae fall within the embryophyte classes, but instead form several monophyla, each containing a rich diversity (Figure 3B). This bolsters the notion that CAD was present in the LCA of zygnematophytes and land plants^52,53,67^. But as the function and structures of CAD homologs vary between the classes, it remains to be experimentally determined which substrates the zygnematophyte CAD homologs accept and under which kinetics they catalyse them. Protein structure predictions hint that streptophyte algal CADs are more like sinapyl alcohol dehydrogenases^49^ (SADs). However, given that (a) *CAD* homologs are differentially regulated in diverse zygnematophytes exposed to stressors^68,69^ and (b) radiated in zygnematophytes, CAD seems to be a potential key player contributing to the chemodiversity of zygnematophytes.

Phosphorelay-based signaling is key in molecular physiological adjustments of land plants. In this response system, histidine kinases homologous to the ARABIDOPSIS HISTIDINE KINASES (AHK) are major players, acting in cytokinin-based signaling^70-72^, their connections to stress response^73^, and frequent responders in transcriptome data of stressed zygnematophytes^69,74-76^. Our maximum likelihood phylogenetic analysis recovers massive expansions in zygnematophyte AHKs in a lineage-dependent manner (Figure 3C). All clades in the gene tree contain proteins with Response regulator receiver domains (Response_reg), His Kinase A (phosphor-acceptor) domains (HisKA) and Histidine kinase-, DNA gyrase B-, and HSP90-like ATPase Histidine Kinase domains (HATPase_c). Yet, we also recover AHK1 and AHK5 as clear orthologous monophyla, whilst there are several co-orthologs to AHK2,3,4. In *Arabidopsis*, AHK2, 3 and 4 bind cytokinin via their extracellular CHASE domain^71,77,78^. Next to the land plant specific clade there are large expansions in zygnematophyte histidine kinases containing CHASE domains. We identified several zygnemtatophyte histidine kinases that contain a PAS fold or domain. These domains are widespread across the tree of life and play a role in light sensing in plants^79^. PAS domain containing histidine kinases that are not phytochromes also exist in *Physcomitrium patens*, other non-seed plants as well as in *Klebsormidium nitens*; in *Physcomitrium* they are regulated by light and play a role in gametophore development^80^. We recovered three of the five *Klebsormidum* PAS-kinases identified by Ryo et al.^80^ in our dataset within the zygnematophyte PAS kinase clade. Our analysis shows that PAS-domain-containing histidine kinases are abundant and likely to be important in zygnematophytes. We also identified histidine kinases with PAS domains in the clade containing AHK5. The majority of the PAS domains correspond to the zygnematophyte sequences within that clade. On balance, Zygnematophyceae have a rich and diversified repertoire of AHKs, likely speaking of their extensive intracellular signaling.

Overall, our data capture the shared occurrence of what has been coined^33^ ‘land plant heritage genes’—genes that constitute a toolkit key for signature traits of land plants. However, this shared molecular toolkit is, in light of loss and divergence, only graspable by using a phylodiverse dataset^12^. With our vastly extended sampling of functional genomic space, several order-dependent patterns of loss and expansion can be gleaned— simultaneously bolstering the need for further whole genome sequencing efforts into zygnematophyte diversity.

### Conclusion: dynamic trait evolution in zygnematophytes

We inferred a robust phylogenomic framework for zygnematophytes, the closest algal relatives of land plants that pays homage to their species richness. Our framework recovers pronounced deep genetic divergences within Zygnematophyceae. This is not only evident from the genetic distances in the tree, but through the order-specific occurrence and within-order radiations, of genes for signature traits, and strong reductive evolutionary forces acting on the genomes^12^. This is mirrored by the diversity in cellular features and growth patterns, including repeated shifts between uni- and multicellular body plans^7,10^, division patterns^81^, plastid morphology^13^, and more. Our data bolster the concept that the LCA of Zygnematophyceae was an ancestral unicell (Figure 2); transitions to multicellularity occurred several times independently, with a few ancient transitions in the LCAs of classical filamentous genera such as *Spirogyra, Zygnema*, and *Mougeotia* (Figure 2). Overall, we infer seven transitions to filamentous body plans, to which broader sampling will likely add more. A comparative and complementing phylodiverse approach that accounts for loss and lability^12^ (some of which might be environmentally inducible^82^) is thus needed to reliably define synapomorphies of the LCA of algae and all land plants.

## Acknowledgement

We thank Prof. T. Friedl from the Department of Experimental Phycology and Culture Collection of Algae (SAG) at the University of Göttingen for his support. We further thank the Culture Collection of Algae at the University of Texas at Austin (UTEX) and ACOI - Coimbra Collection of Algae for providing algal strains. J.d.V. is grateful for funding by the German Research Foundation grant 509535047 (VR 132/10-1) and the grants 440231723 (VR 132/4-1), 528076711 (VR 132/13-1) within the framework of the Priority Programme “MAdLand – Molecular Adaptation to Land: Plant Evolution to Change” (SPP 2237). J.d.V. further thanks the European Research Council for funding under the European Union’s Horizon 2020 research and innovation programme (Grant Agreement No. 852725; ERC-StG “TerreStriAL”) and the Horizon Europe programme (Grant Agreement No. 101230161; ERC-CoG “StreptoProgram”). I.I. is grateful for funding to the Spanish State Research Agency (AEI) MICIU/AEI/10.13039/501100011033, ESF+ and ERDF/EU (Grants RyC2022-038245-I and PID2023-152168NB-I00) and the Spanish Research Council (Ayudas Atracción de Talento CSIC PIE Grant 20243AT022). M.J.B, C.F.K and E.S.G. are grateful for support through the IMPRS Genome Science

## STAR METHODS

### RESOURCE TABLE

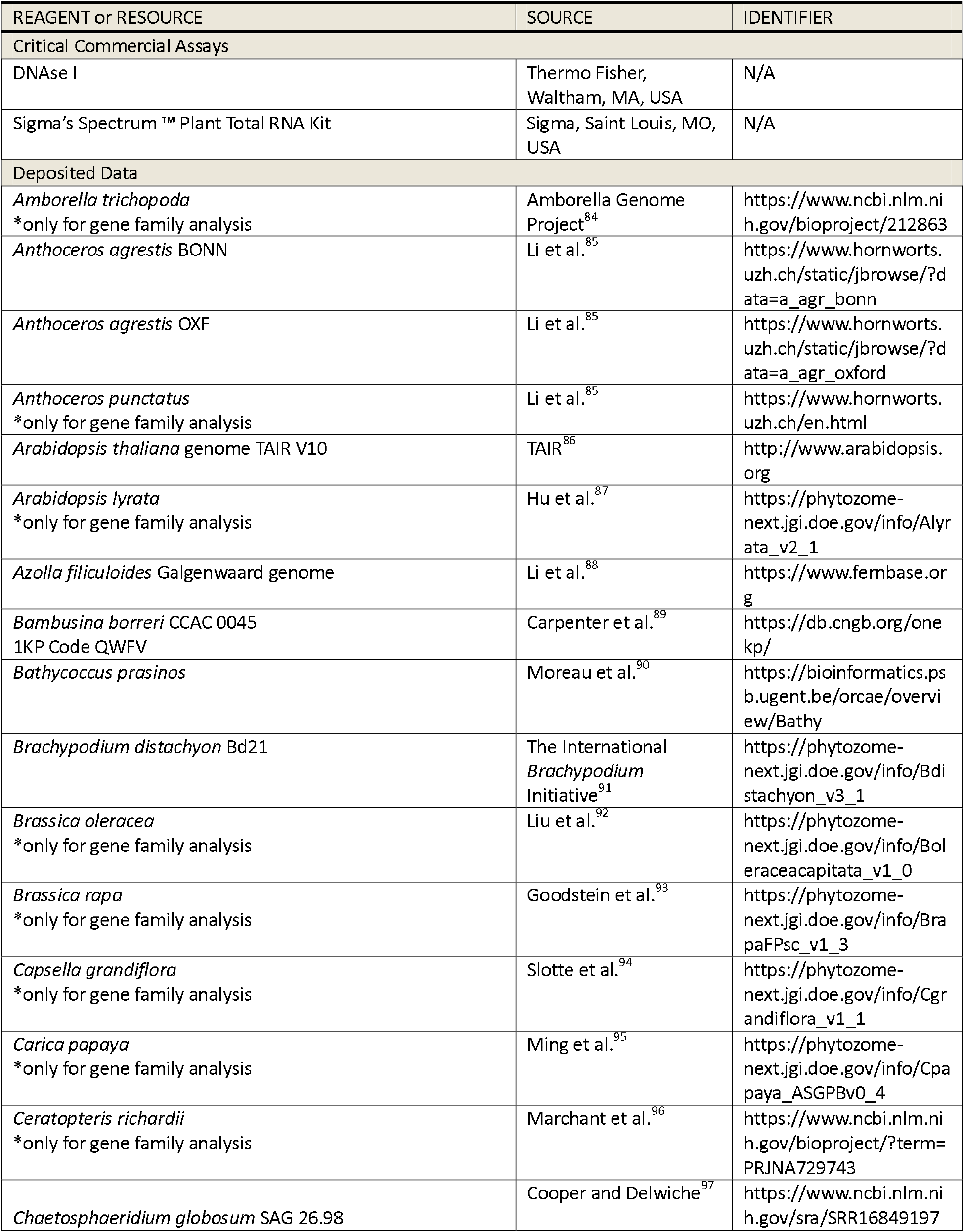
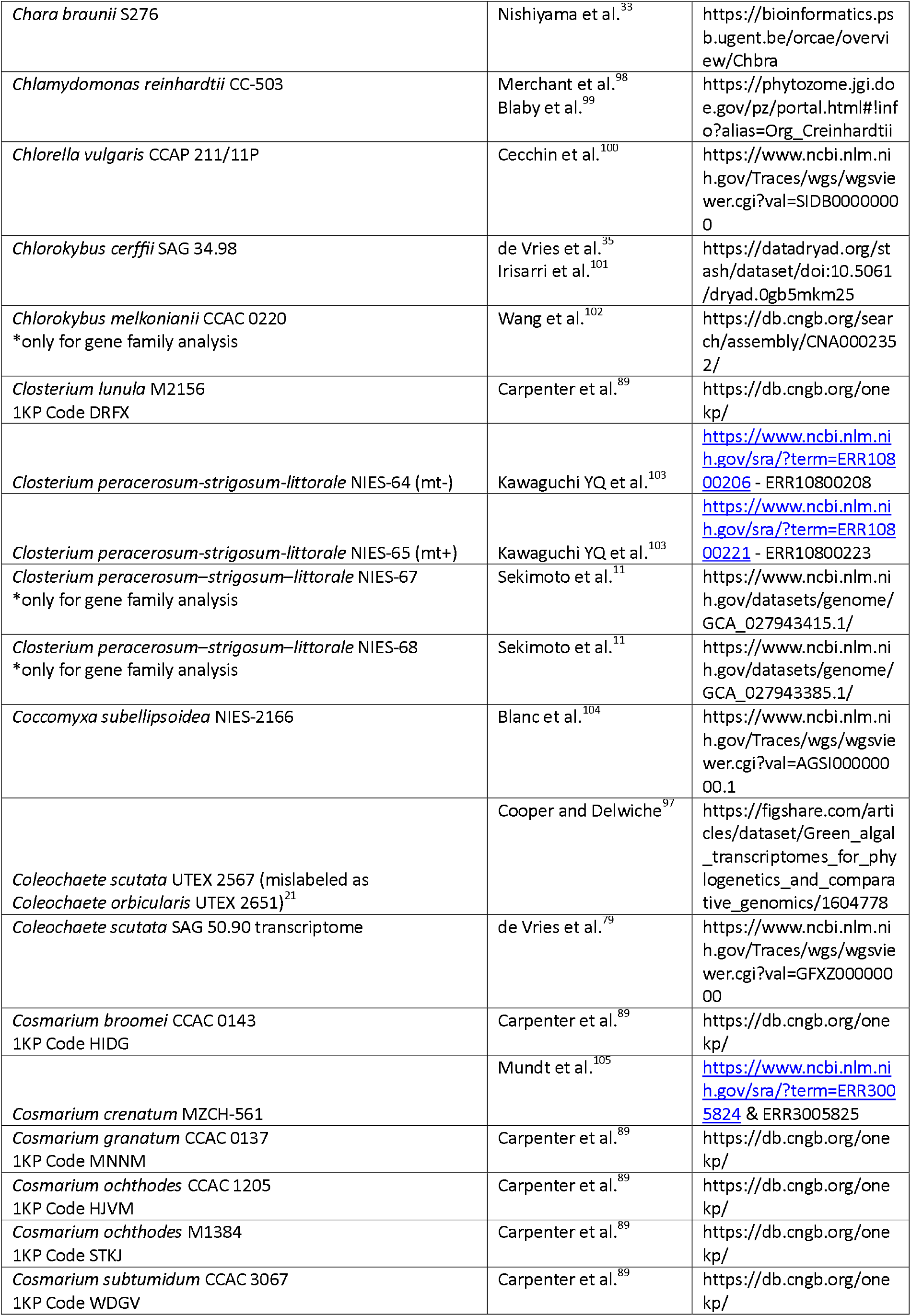
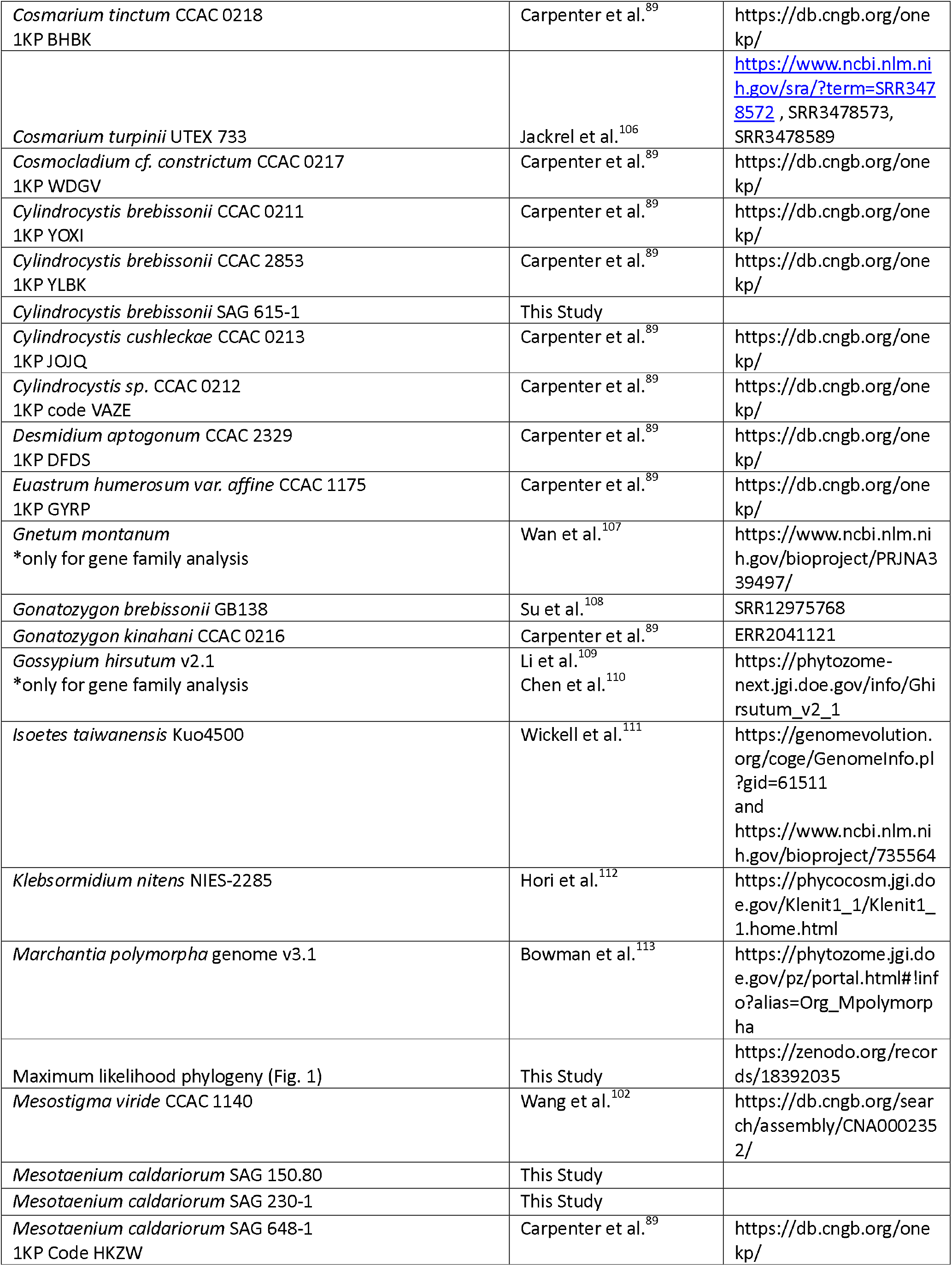
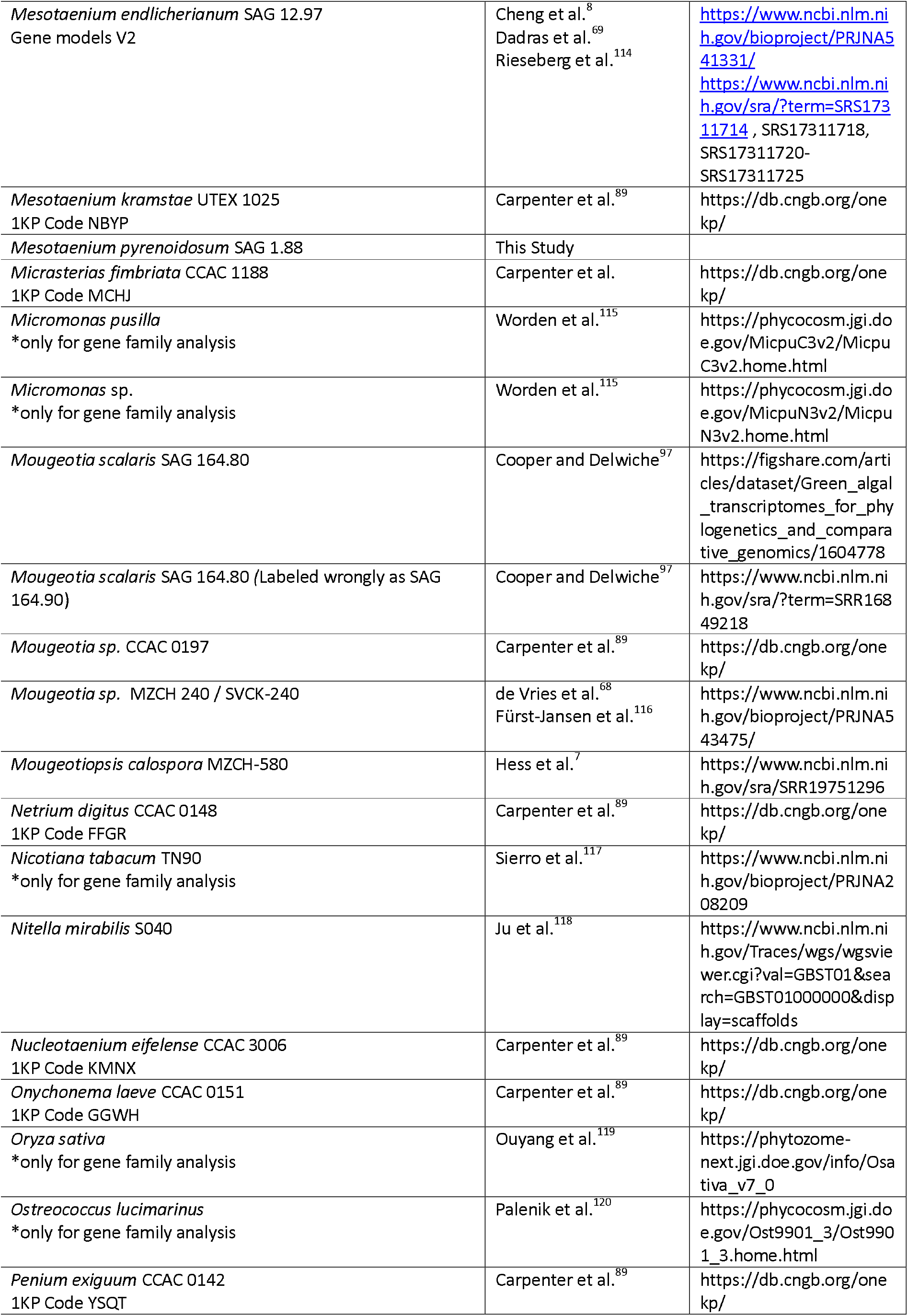
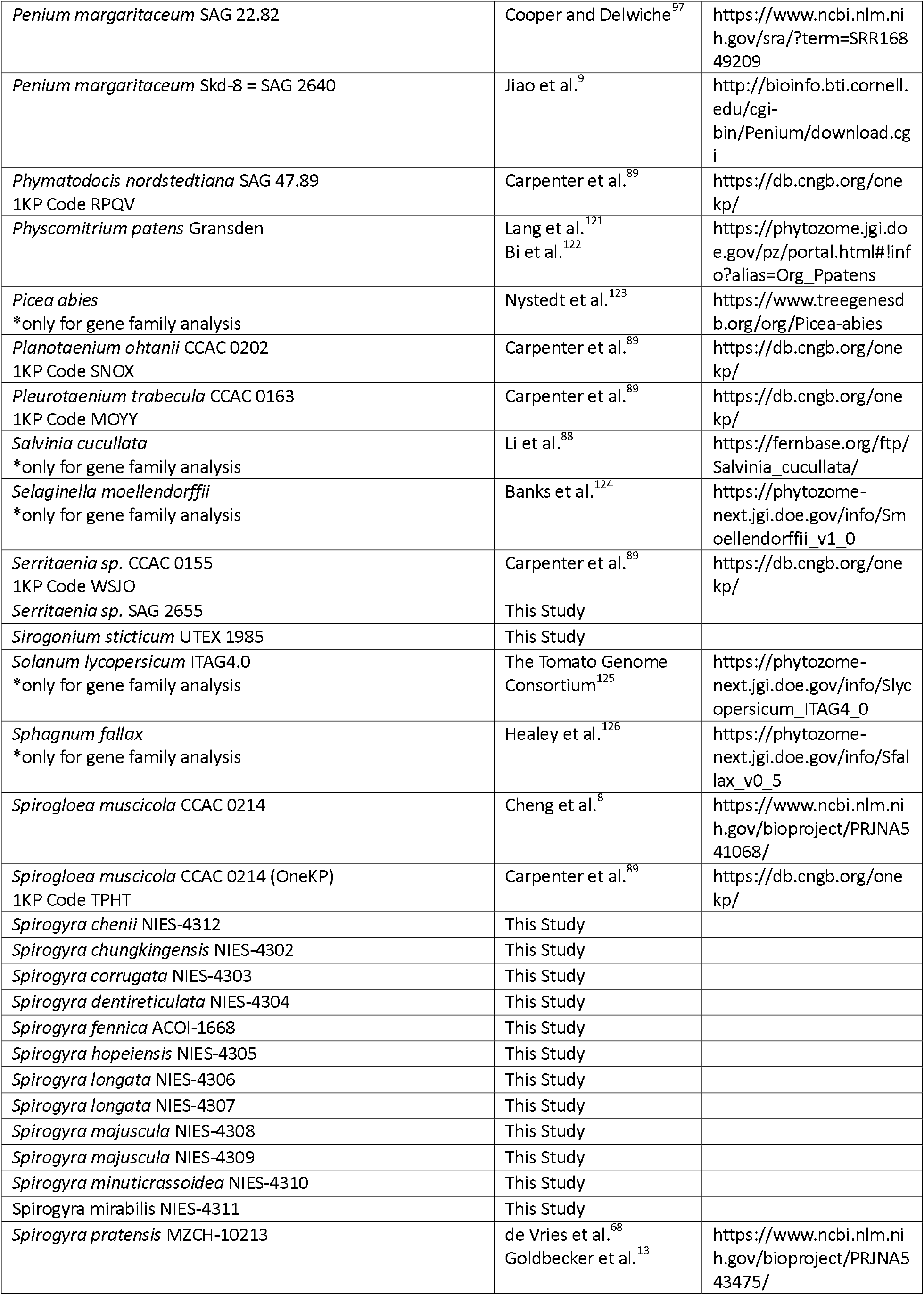
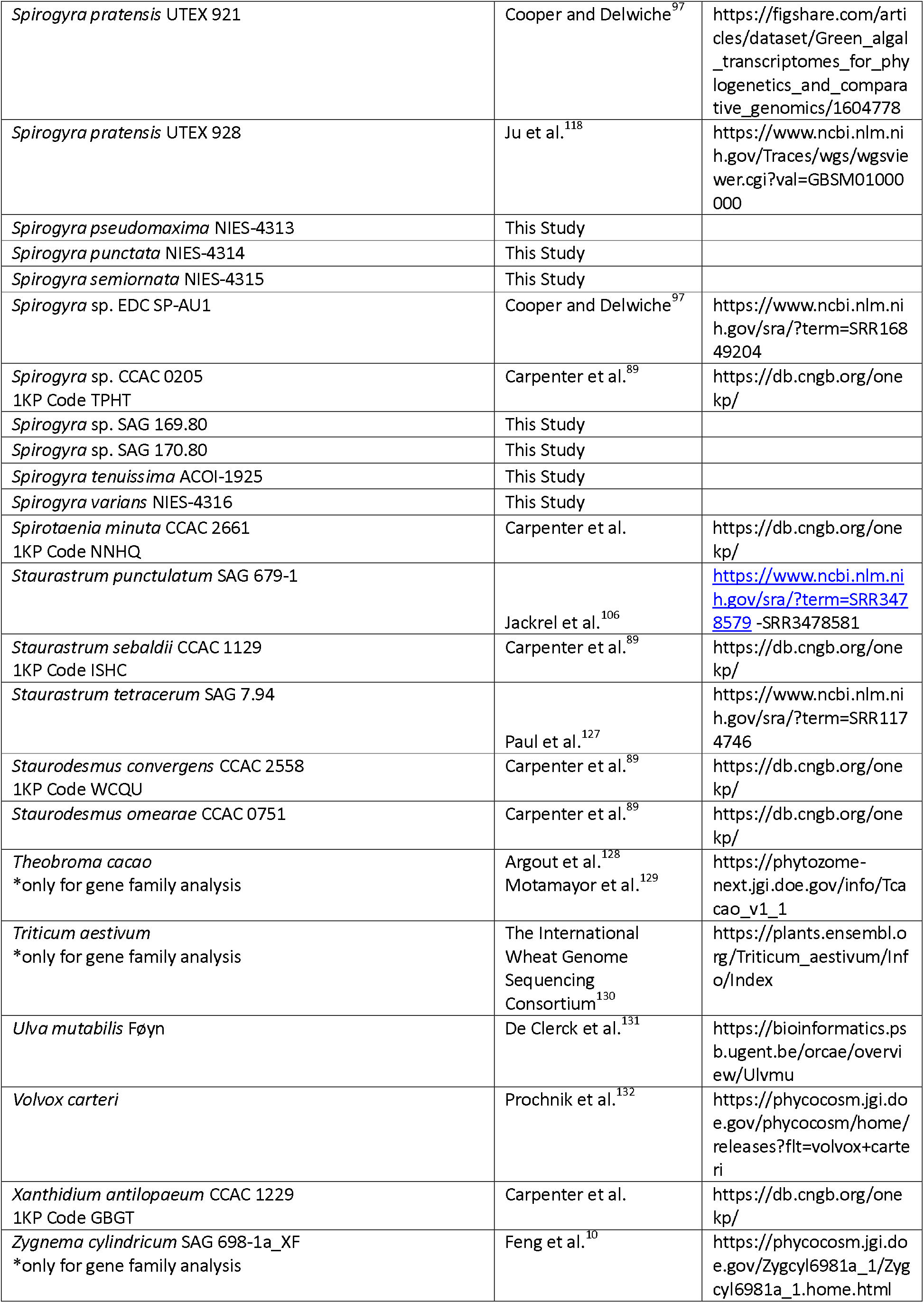
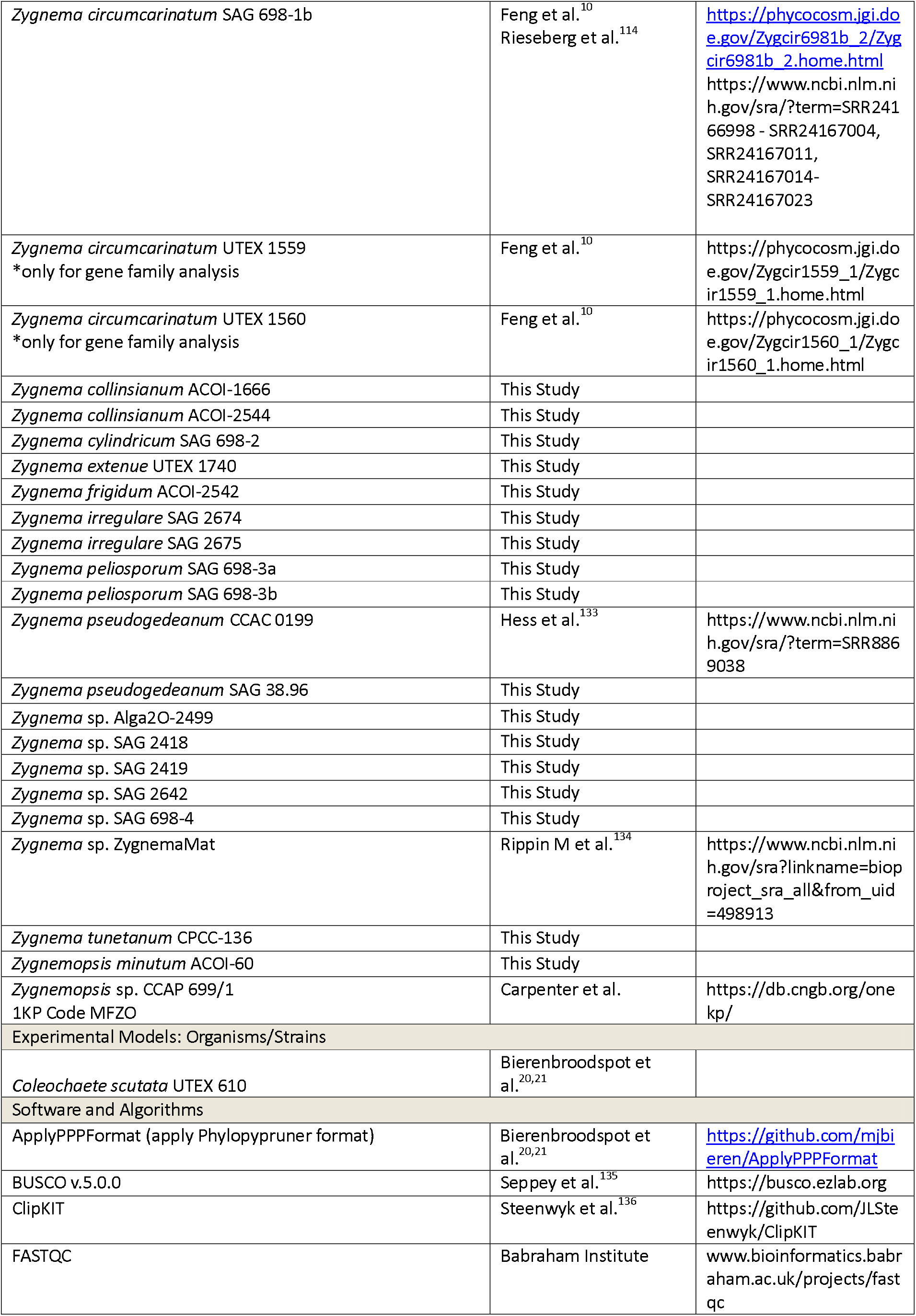
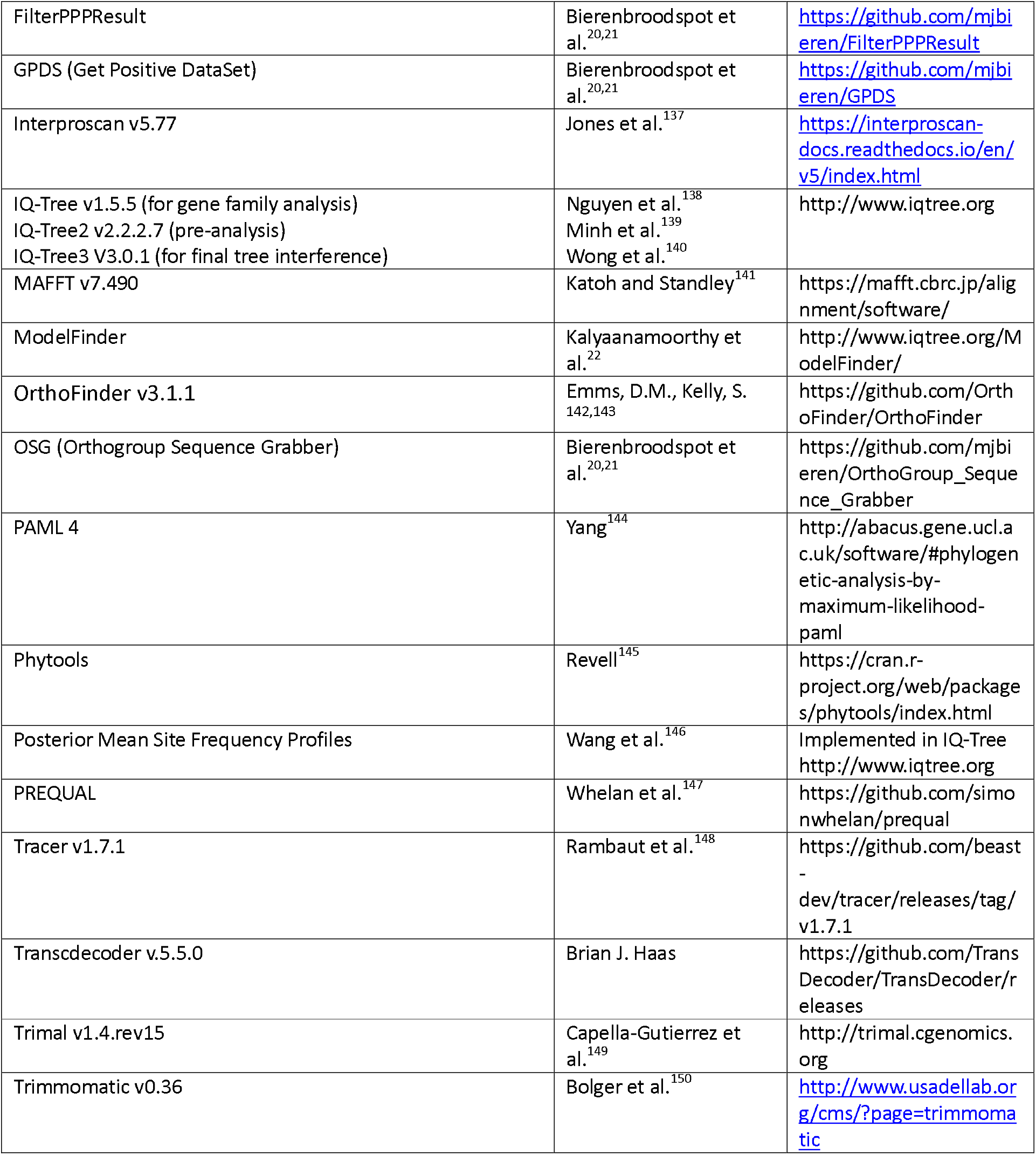

## RESOURCE AVAILABILITY

### Lead contact

Further information and requests for resources and reagents should be directed to and will be fulfilled by the lead contact, Jan de Vries.

### Materials availability

This study did not generate new unique reagents.

### Data and code availability

- All RNA-seq data generated in this study have been deposited in the NCBI
- All resulting data and analytical outputs are accessible under https://doi.org/10.5281/zenodo.18392035 on Zenodo.
- A comprehensive code and methodological overview is available at https://github.com/mjbieren/Zygnematophyceae_Phylogenomics

## EXPERIMENTAL MODEL AND SUBJECT DETAILS

### Algal strains

Strains were obtained from the Culture Collection of Algae at Göttingen University (SAG)^151^, the microbial culture collection of the National Institute for Environmental Studies in Japan (NIES), the Coimbra Collection of Algae (ACOI), the Microalgae and Zygnematophyceae Collection Hamburg (MZCH), and the culture collection of algae at the University of Texas at Austin (UTEX).

## METHOD DETAILS

### Algal culturing

All cultures were maintained for six weeks in 3NBBM or MiEB12 liquid medium (medium 26a in Schlösser^152^, and medium 7 in Schlösser^153^) under standardized laboratory conditions (18⍰°C; 14:10 h light:dark cycle; 25–35 μmol photons m^−2^ s^−1^) using full-spectrum fluorescent lighting.

### Light microscopy

High-resolution images of the studied strains were obtained with an Olympus BX-60 microscope (Olympus, Japan) with DIC equipped with a ProgRes C14plus camera and the ProgRes CapturePro Software (version 2.9.01) (JENOPTIK AG, Jena, Germany). All investigated strains were examined in week six of cultivation.

### RNA extraction

For the RNA extraction of the strains, 50 mL of six-weeks-old liquid culture were centrifuged for 5 min at 20° C and 11000 rpm and the supernatant was removed. Pellets were transferred into 1.5ml BioMasherII tubes (Nippi, Japan) and each sample was disrupted using PowerMasherII (Nippi, Japan) after freezing in liquid nitrogen. RNA extraction was done using the Spectrum Plant Total RNA Kit (Sigma-Aldrich Chemie GmbH, Germany) according to the manufacturer’s instructions. DNAse I treatment (Thermo Fisher, Waltham, MA, USA) was applied to the RNA samples, and their integrity and concentration were assessed using a 1% agarose gel with a Midora green stain, and Nanodrop (Thermo Fisher), respectively. The RNA samples were shipped on dry ice to Novogene (Munich, Germany).

### RNA sequencing, transcriptome assembly, and transcriptome data

At Novogene (Munich, Germany), RNA quality was verified using a Bioanalyzer (Agilent Technologies, USA). Libraries were prepared using poly-A enrichment and directional mRNA library construction protocols. After the libraries were quality checked, sequencing was conducted on an Illumina NovaSeq 6000 platform using dual-indexed Novogene adapters: 5⍰-AGATCGGAAGAGCGTCGTGTAGGGAAAGAGTGTAGATCTCGGTGGTCGCCGTATCATT-3⍰ for read 1 and 5⍰-GATCGGAAGAGCACACGTCTGAACTCCAGTCACGGATGACTATCTCGTATGCCGTCTTCTGCTTG-3⍰. We conducted transcriptome sequencing on 43 isolates belonging to the different order of the Zygnematophyceae. All transcriptomes were assembled *de novo* with Trinity^154^ v2.15.1, following adapter trimming with Trimmomatic^150^ (--trimmomatic “ILLUMINACLIP:novogene_adapter_sequences.fa:2:30:10:2:keepBothReads LEADING:3 TRAILING:3 MINLEN:36⍰) and default settings. Splicing isoforms were collapsed into SuperTranscripts^155^ using Trinity’s implementation. Completeness was assessed with BUSCO^135^ v5.4.3 using the ‘eukaryota_odb10’ dataset, yielding an average completeness of 94.84%. Protein-coding sequences were predicted with TransDecoder v5.5.0, retaining only the longest open reading frame per transcript (--single_best_only).

### Phylogenomic analysis: preamble

Phylogenomic analyses and ancestral character state reconstruction were conducted following the framework outlined in Bierenbroodspot et al.^20^, with the following methodological refinements. For orthogroup selection, we utilized only the Phylogenetic Hierarchical Orthogroups at the tree root (N0.tsv file produced by OrthoFinder v3.1.1^142,143^) in conjunction with Orthogroup Sequence Grabber (OSG)^20,21^. A streamlined version of the COGS software^20,21^ was implemented, excluding one redundant MAFFT^141^/IQ-TREE^138-140^ alignment step. Similarly, the final ClipKit^136^ trimming step was omitted, as it did not enhance alignment quality under the revised workflow.

### Phylogenomic analysis: dataset construction for phylotranscriptomics

To eliminate contamination from *de novo* transcriptome assemblies, sequence similarity searches were conducted using the MMseqs2 tool^156^ against a custom database containing proteins from the axenic strains MZCH-10213, SAG 1.88, SAG 12.97, SAG 615-1, SAG 698-1a, and SAG 698-1b as positive control, and potential contaminants including RefSeq representative genomes of bacteria (43,588), fungi (1,225), invertebrate animals (7,213), virus (13,134), archaea (1,120), protists (893), mitochondrial genes (240,117), plasmids (4,710), and plastids (12,307) (downloaded July 11, 2023). An iterative approach was used for MMseqs2 (--start-sens 1 --sens-steps 3 -s 7 --alignment-mode 3 --max-seqs 10), and decontamination was completed with GPDS^20,21^. Only sequences whose best match was against the combined positive set were retained. Potential contaminant-matching sequences were archived and are available via Zenodo https://doi.org/10.5281/zenodo.18392035

### Phylogenomic analysis: phylotranscriptomic analysis

Orthogroup inference was performed using OrthoFinder^143^ v3.1.1 with a species tree following the framework of the One Thousand Plant Transcriptomes Initiative^2^, including chlorophytes, *Chlorokybus cerffii* SAG 34.98, *Mesostigma viride* NIES-296, and *Klebsormidium nitens* NIES-2285, alongside multiple Phragmoplastophyta and Coleochaetophyceae (see “09_Orthofinder.zip” on Zenodo). A total of 60,932 orthogroups were identified. Two orthogroup subsets were generated using OSG^20,21^: one enriched for ingroup taxa (13,409 orthogroups, filter: 2 of 4 groups) and another for outgroups (7,253 orthogroups, filter: 10 of 17 groups). All data are available at Zenodo (“10_OSG.zip”).

Multiple sequence alignments were performed with MAFFT^141^ v7.304 under default settings. Maximum likelihood trees were constructed using the IQ-TREE 2 software^139^ v2.2.2.7 with automated model selection via ModelFinder^22^ (selection of best-fit models was based on Bayesian Information Criterion), using SH-like aLRT (-alrt 1000) and ultrafast bootstrap replicates (-bb 1000). Tree files were reformatted for PhyloPyPruner v1.2.4 using ApplyPPPFormat (https://github.com/mjbieren/ApplyPPPFormat). PhyloPyPruner (https://github.com/fethalen/phylopypruner) was used to remove paralogs. For the orthogroups subset biased towards the ingroup we used the following settings: --mask pdist --prune MI --min-taxa 3 --trim-lb 5 --min-support 0.75 --threads 50 --trim-divergent 1.25 -- jackknife, yielding 93,280 ortholog alignments. For the outgroup-centered subset, we used: - -min-taxa 10 --min-gene-occupancy 0.1, yielding 5,960ortholog alignments. After taxonomic filtering to ensure a balanced taxonomic representation in alignments (FilterPPPResult, -t 10 (of 110)), we retained 17,588 ingroup-rich and (FilterPPPResult, -t 2) 5,923 outgroup-rich loci. These were merged using COGS^21^ and then pruned to 5,225 unique orthogroups. A final PhyloPyPruner run produced 80,973 loci; after filtering (FilterPPPResult, -t 30), 2,243 loci remained. These were masked using PREQUAL v1.02 and re-aligned with MAFFT (G-INS-i, ‘-- allowshift --unalignlevel 0.8’). Gappy columns (>75% gaps) were removed using ClipKIT^136^ v2.0.1. The final concatenated matrix was created using phyx^157^ comprising 126 taxa, 2,243 loci, and 579,537 aligned amino acid sites. Phylogenies were reconstructed with IQ-TREE 3 V3.0.1 under the LG+C60+G model, using 1000 SH-aLRT and UFBoot^158^ replicates.

### Ancestral Character State Reconstruction

Ancestral state reconstruction (ACSR) was conducted using Phytools^145^, employing Yang’s re-rooting method^159^. Traits were coded with a 5-state model: (1) unicellular, (2) multicellular, (3) Package-like, (4) Bona fide filaments, and (5) Chain-like filaments. Analyses assumed symmetric transition rates between unordered states.

### Gene family analysis

For gene family analysis, we worked with a dataset of predicted proteins derived from (i) the transcriptomes generated in this study and two available ones and (ii) published genomes of phylodiverse Chloroplastida (see Resource Table). We searched for homologs of the genes of interest using representative sequences from *Arabidopsis* as a query in a BLASTp search against the full protein dataset. An e-value cutoff of 10^-7^ was applied; in case of large gene families, the number of species was reduced. All sequences were aligned using MAFFT^141^ v7.490 (L-INS-i) and a maximum likelihood phylogeny was computed using IQ-TREE^138^ with 1000 UFBoot^160^ pseudoreplicates and best-fit models chosen according to Bayesian Information Criterion by ModelFinder^22^. The AHK gene family tree was annotated using PFAM domains predicted by interproscan^137^ v5.77-108.0. The domains were consolidated per clade. A clade was annotated to contain a domain when it appeared in several proteins within that clade. Outlier domains, e.g. only predicted for 1 protein within the whole clade, were ignored. We summarized PFAM PAS fold and PAS domain annotations (PF00989, PF08447, PF08448, PF13188, PF13426) with one symbol.

## QUANTIFICATION AND STATISTICAL ANALYSIS

The final maximum likelihood phylogenomic tree was inferred using the LG+C60 model with 1000 SH-aLRT and 1000 ultrafast bootstrap replicates. Additional analyses incorporating site-heterogeneous PMSF models were conducted using a guide tree inferred with LG+C60+F+Γ. These consistently recovered the same topology as the original LG+C60-based tree.

ModelFinder^22^ identified LG+C60 as the best -fitting model for protein evolution compared to the more simplified mixture model LG+I+G4, as indicated by improved likelihood scores and a lower Bayesian Information Criterion (LG+C60: Log(Likelihood) = −15,486,259.27; Bayesian Information Criterion = 30,975,836.031; LG+I+G4: Log(Likelihood) = −15,683,898.390; Bayesian Information Criterion = 31,371,127.541). Themaximum likelihood phylogenomic tree was inferred under the LG+C60 model using 1,000 SH-aLRT and 1,000 ultrafast bootstrap replicates. To account for site heterogeneity, an additional analysis was performed using the PMSF approximation with LG+C60+F+Γ, guided by the previously inferred tree. This analysis consistently recovered the same tree topology, reinforcing the robustness of the phylogenetic inference across model frameworks.

For gene family analyses, 144 protein models were tested using ModelFinder^22^ and Bayesian Information Criterion was used to determine the best-fit model. 1000 UFBoot^160^ pseudoreplicates were computed per gene tree.

