## Supplemental Figure S1 for "A deep genetic structure phylogenomically frames the closest algal relatives of land plants"

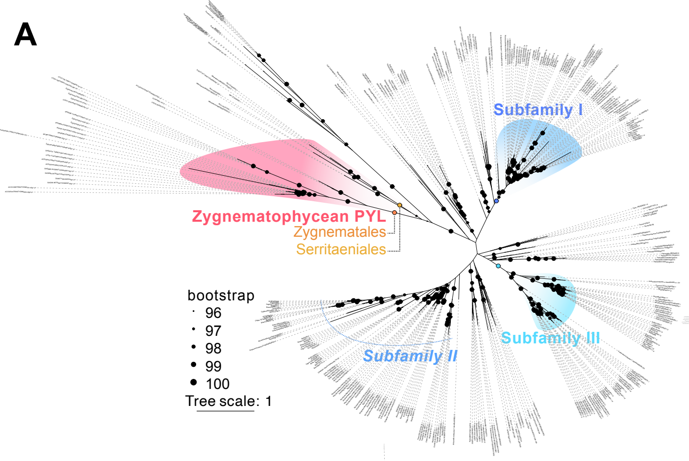


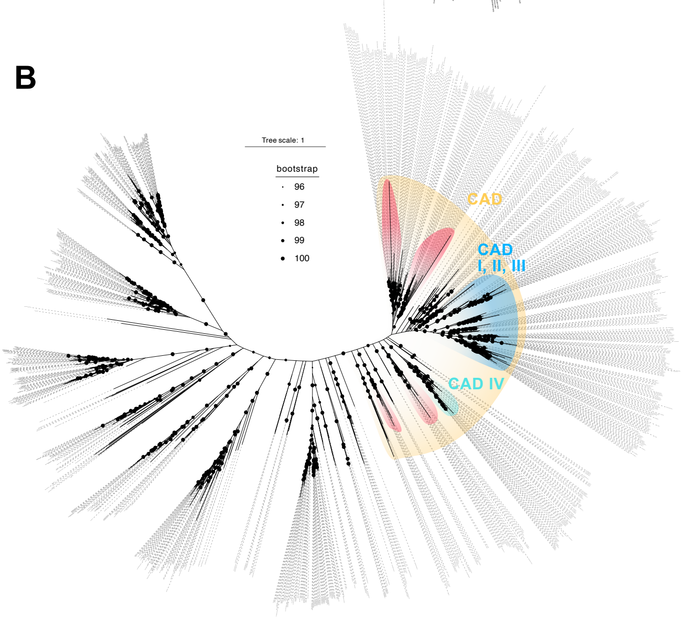


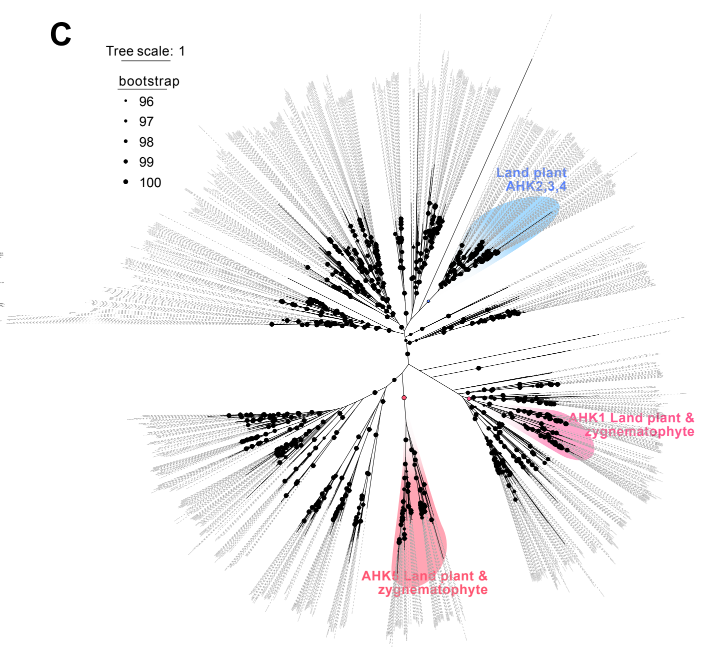


**Figure S1. Fully labelled phylogenetic trees of signature gene families. Related to Figure 3.** Fully labeled phylogenies shown in Figure 3.
